# Enzymatic analysis of yeast cell wall-resident GAPDH and its secretion

**DOI:** 10.1101/2020.10.12.336925

**Authors:** Michael J. Cohen, Brianne Philippe, Peter N. Lipke

## Abstract

In yeast, many proteins are found both in the cytoplasmic and extracellular compartments, and consequently it can be difficult to distinguish non-conventional secretion from cellular leakage. We therefore monitored extracellular glyceraldehyde-3-phosphate dehydrogenase (GAPDH) activity of intact cells as a specific marker for non-conventional secretion. Extracellular GAPDH activity was proportional to the number of cells assayed, increased with incubation time, and was dependent on added substrates. Preincubation of intact cells with 100μM dithiothreitol increased the reaction rate, consistent with increased access of the enzyme after reduction of cell wall disulfide crosslinks. Such treatment did not increase cell permeability to propidium iodide, in contrast to effects of higher concentrations of reducing agents. An amine-specific membrane-impermeant biotinylation reagent specifically inactivated extracellular GAPDH. The enzyme was secreted again after a 30-60-minute lag following the inactivation, and there was no concomitant increase in propidium iodide staining. There were about 4 × 10^4^ active GAPDH molecules per cell at steady state, and secretion studies showed replenishment to that level one hour after inactivation. These results establish conditions for specific quantitative assays of cell wall proteins in the absence of cytoplasmic leakage and for subsequent quantification of secretion rates in intact cells.

**Importance:** Eukaryotic cells secrete many proteins, including many proteins that do not follow the classical secretion pathway. Among these, the glycolytic enzyme glyceraldehyde-3-phosphate dehydrogenase (GAPDH) is unexpectedly found in the walls of yeasts and other fungi, and in extracellular space in mammalian cell cultures. It is difficult to quantify extracellular GAPDH, because leakage of just a little of the very large amount of cytoplasmic enzyme can invalidate the determinations. We used enzymatic assays of intact cells, while also maintaining membrane integrity. The results lead to estimates of the amount of extracellular enzyme, and its rate of secretion to the wall in intact cells. Therefore, enzyme assays under controlled conditions can be used to investigate non-conventional secretion more generally.

## Introduction

The glycolytic enzyme glyceraldehyde-3-phosphate dehydrogenase (GAPDH) is unexpectedly found in the walls of *Saccharomyces cerevisiae, Candida albicans*, and *Paracoccidiodides brasiliensis* (1–8). The enzyme is also secreted from mammalian cells in culture (9–11). Like many glycolytic proteins, GAPDH is a moonlighting protein with additional roles both within the cell (12, 13) and externally; in *C. albicans* cell wall GAPDH binds fibronectin (14), and in *S. cerevisiae* its secreted form is cleaved into antimicrobial peptides (15, 16). Recent cell wall proteomics work has shown cell wall localization of GAPDH in *C. albicans* and non-albicans species (17, 18). The protein is present in walls, and in many cases its concentration is increased after growth in media that mimic mammalian conditions. Thus GAPDH is a common wall marker in pathogenic yeasts and may be important in host-pathogen interactions.

All three isoforms of GAPDH, encoded by *TDH1, TDH2*, and *TDH3*, are enzymatically active in *S. cerevisiae* walls (1), but cell surface quantities and pathways leading to secretion remain elusive. Mass spectrometry of protease-treated cell walls or of intact cells generates peptides from enolase, alcohol dehydrogenase, and GAPDH, as well as cytosolic chaperones such as Ssa1, and Ssa2 in both *S. cerevisiae* (4) and *C. albicans* (6, 19). Thus, GAPDH is prototypical of many unconventionally secreted proteins (20), as defined by their presence in the extracellular compartment despite their lack of canonical secretion signal peptides.

*S. cerevisiae* can be engineered to display and anchor enzymes on the cell wall for biofuel production (21), bioremediation (22), or library screening (23). The cell walls consist of polysaccharides including β1,3 glucans, β1,6 glucans, and chitin, and a large number of proteins. These cell wall resident proteins crosslink the saccharides, act as adhesins, regulate metabolic activities, and perform other functions (24–26). Most of these proteins are secreted through the conventional secretion-signal-dependent pathway that processes the proteins through the ER and Golgi (27, 28). This pathway was famously elucidated through a combination of enzymology and genetic screens (28). Temperature sensitive *S. cerevisiae* secretory mutants were generated, and at non-permissive temperatures they showed defects in invertase and acid phosphatase secretion (29).

Yeast have also been used to study unconventional protein secretion of proteins which lack a signal peptide. *S. cerevisiae* expressing the mammalian protein Galectin-1 could secrete it without using its classical secretory system (30), much as the protein behaves in mammalian cells (31). Mutational studies in in *S. cerevisiae* identified an Acb1 secretory mechanism that requires autophagy, Golgi proteins, as well as endosome components (32, 33). The chitinase Cts1 from *Ustilago maydis* was used to study a novel form of unconventional protein secretion at budding sites (34). Yeast species are also used to characterize unconventional secretion secretion into extracellular vesicles (20, 35–37). Thus, yeast is now a classic model for study of secretory pathways in general.

We are interested in studying unconventional secretion of proteins such as GAPDH using an enzymology approach. However, GAPDH is abundant in cytosol, there is a critical need to obtain cell wall extracts while avoiding cytosolic contamination. We therefore describe procedures for quantitative assay of extracellular GAPDH, and techniques for its extraction without contamination by cytosolic enzyme.

## Results

### GAPDH activity in the wall of intact *S. cerevisiae*

We verified that GAPDH is enzymatically active in the wall of *S. cerevisiae* strain BY4743 by resuspending cells in 1 mM NAD, 1mM glyceraldehyde-3-phosphate, 100 μM DTT, and triethanolamine phosphate (TEA) buffer (pH 8.6) in a 200 μL reaction, using methods similar to that of Delgado et al (1). However, we extended their results by establishing the wall-associated activity on a per cell basis. We suspended different concentrations of *S. cerevisiae* in GAPDH substrates for 30 minutes then measured NADH production. Yeast cells were pelleted by centrifugation, and NADH was measured as A_340_. NADH production was linear with cell number up to 1.5 x 10^6^ per 200 μL reaction (Fig. 1A). Therefore, subsequent experiments used a maximum of 1x 10^6^ cells in 200 uL, with a majority of trials using 5 x 10^5^ cells. The reaction rate was linear between 30 and 60 minutes, but showed a lag before that time (Fig. 1B). The origin of the lag is addressed in the next section.

**Fig. 1.**
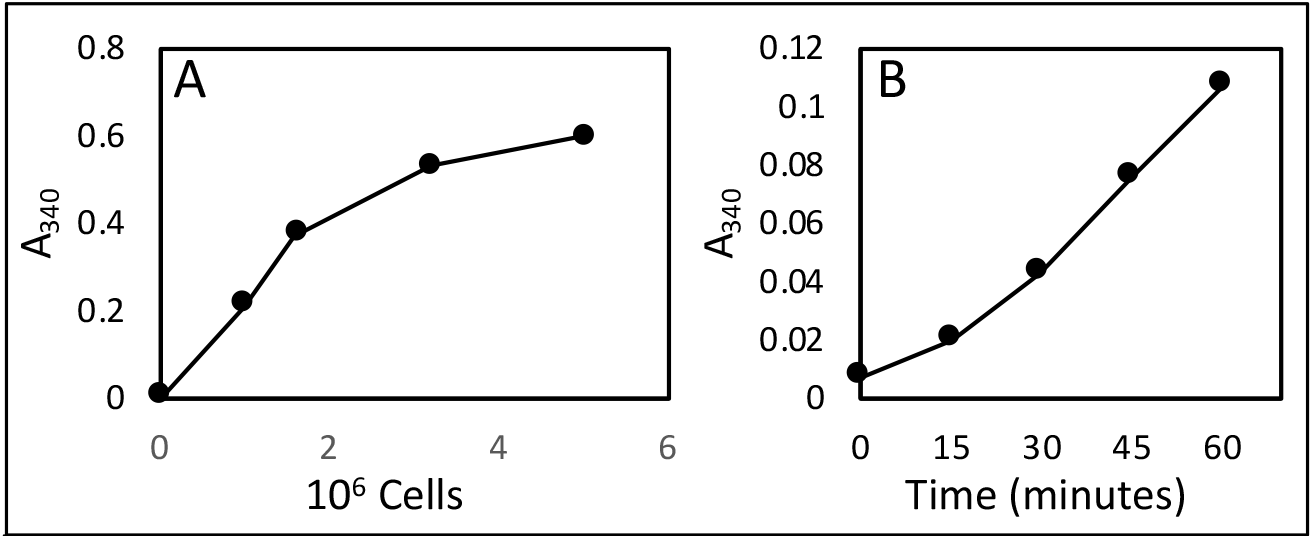
Enzymological characteristics of cell wall GAPDH assays. A) Dependence on cell number in a 30 min assay. B) Time course of the assay with 5 × 10^5^ cells.

Other enzymological controls were as expected. There was no measurable activity in the absence of added NAD+ or glyceraldehyde-3-phosphate. This data shows that all assayed NAD+ reductase activity was due to GAPDH, and there was no endogenous activity due to leakage of either substrate from the cytoplasm. The absorbance spectrum of the product matched NADH, and the optimum reaction pH was 8.6, consistent with known GAPDH properties (38) This pH value suggests that cell surface GAPDH is not enzymatically active during yeast growth, which normally occurs under acidic conditions.

### Cell surface GAPDH activity increases during assays

In 60- and 90-minute assays, the rate of NADH production increased (Fig. 2). This increase emphasized the lag time apparent in Fig. 1B. This result suggested that either GAPDH was accumulating at the surface, or that more extracellular GAPDH became active during the extended incubation in assay buffer.

**Figure 2:**
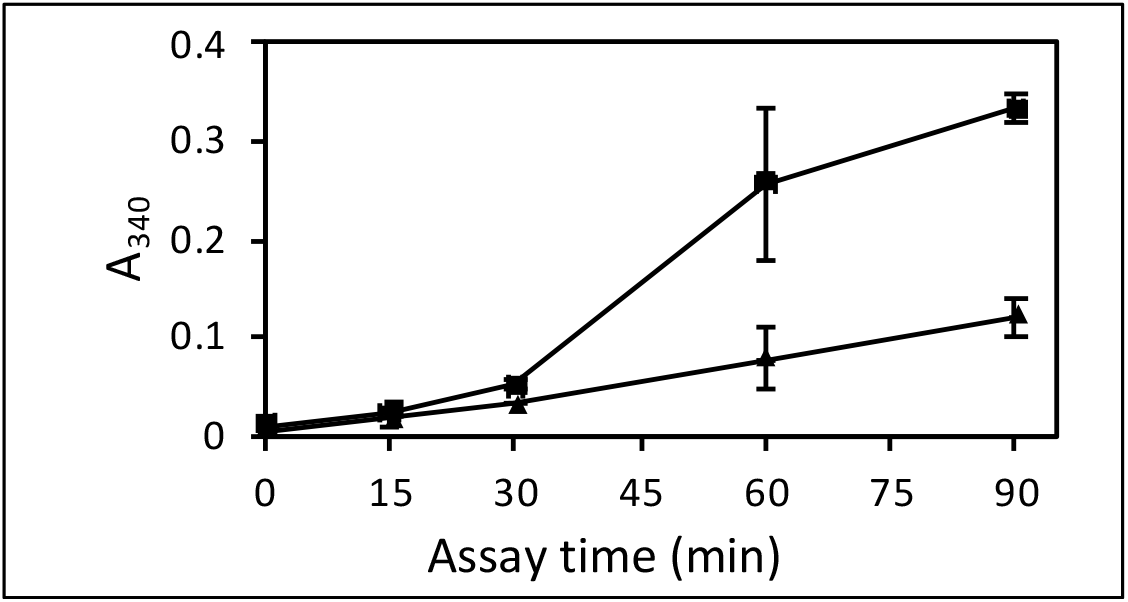
Cell surface GAPDH activity on the presence and absence of DTT. GAPDH assays in the absence (▲) and presence (■) of 100 μM DTT.

GAPDH can be partially oxidized *in vivo*, reducing its activity by 10%, so standard assays contain dithiothreitol (DTT) to keep the enzyme reduced (39). Our assay buffer contained 100 μM DTT, as in Delgado et al. (1, 2). However, DTT can also break disulfide bonds in the wall and increase cell wall porosity (5, 40), exposing more enzyme to substrates (41–43). To test whether DTT was facilitating increased cell wall GAPDH activity, we assayed activity in the presence or absence of 100 μM DTT. The rate of NADH production was greater in the presence of DTT than without it, and the rate of increase was maximum between 30 and 60 min of incubation (Fig. 2). This finding was confirmed in a preincubation experiment. Cells were pre-incubated in assay buffer in the absence of substrate, and in the presence or absence of DTT, 100 μM. Substrates were then added and enzyme activity monitored in standard 30 min assays. Preincubation in DTT increased GAPDH activity at the cell surface in a subsequent GAPDH assay. Therefore, DTT treatment either increased the fraction of surface GAPDH that was enzymatically active, or promoted surface accumulation of the enzyme, or both.

### GAPDH on the surface can be attenuated with a membrane-impermeant covalent modifier

To distinguish between GAPDH already in the wall and newly secreted GAPDH, we took advantage of a membrane-impermeant modifier to deactivate cell-wall associated GAPDH. Attempts to label and extract cell surface GAPDH led to the observation that the biotinylation reagent sulfosuccinimidyl 6-(biotinamido)-hexanoate (Sulfo-NHS-LC biotin) decreased GAPDH activity dramatically, both in cytoplasmic extracts and for the enzyme assayed on the surface of intact cells (Fig. 3). Because Sulfo-NHS-LC biotin is membrane impermeant (44) and will only react with proteins external to the plasma membrane, it can specifically deactivate cell wall GAPDH and leave cytosolic GAPDH unaffected. We treated intact cells with Sulfo-NHS-LC biotin, then washed the cells to remove remaining Sulfo-NHS-LC biotin, resuspended in assay buffer, and assayed for 15-90 min. Unlike the control untreated cells, biotinylated yeast did not have detectable GAPDH activity on their surface for the first 30 minutes (Fig. 3). However, biotinylated yeast showed increasing surface GAPDH between 30 and 90 min (Fig. 3B). This result was consistent with GAPDH being released to the surface from a cellular pool inaccessible to sulfo-NHS-LC biotin, presumably in the cytoplasm. However, the rate of increase of cell surface activity was less than that of the cells not treated with biotinylation reagent. Therefore, about half of the increase shown in Figs. 2 and 3 may be due to secretion of active enzyme to the cell surface during the incubation, but some of the increase may represent more activity of the resident assayable cell surface enzyme, presumably due to the wall permeabilization by DTT.

**Fig. 3.**
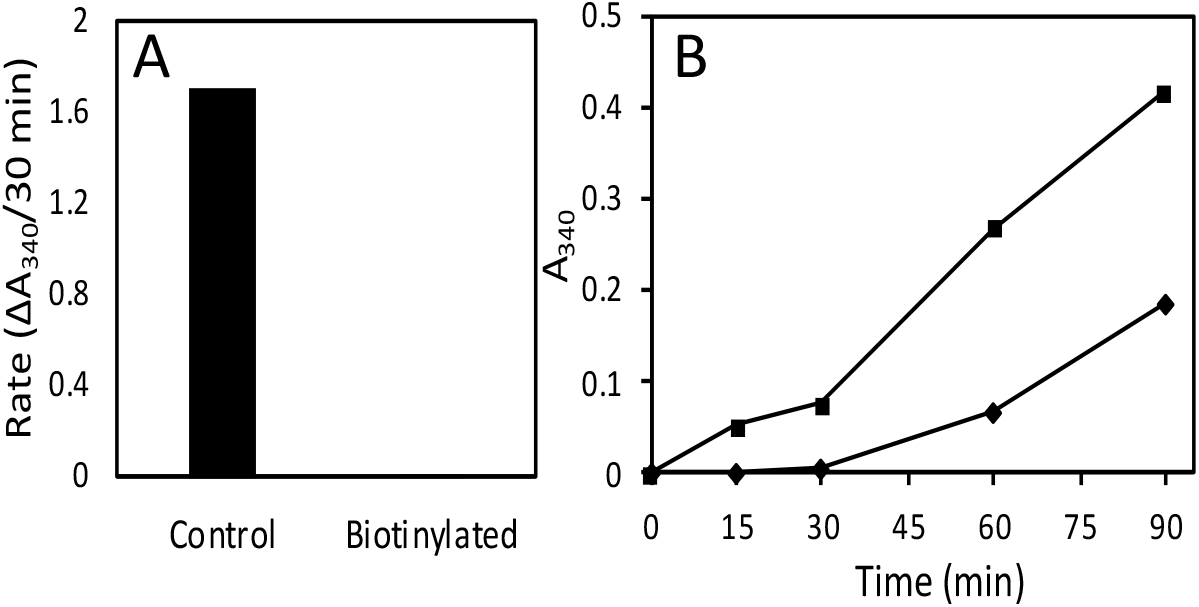
Effect of sulfo-NHS-LC biotin on GAPDH activity. **A)** Cytosolic Lysate biotinylated with 1mg/mL Sulfo-NHS Biotin has low GAPDH activity. Lysate was biotinylated, 10 μL of 1:5 dilutions of lysate was loaded onto a microtiter plate, 90 μL of substrates were added and A_340_ was monitored over 30 minutes. **B)** Yeast grown to an OD of 0.7 were biotinylated (♦) for 1 hr in PBS pH 7 with 1mg/mL sulfo-NHS-LC biotin or PBS (■), and then suspended in TEA buffer containing GAPDH substrates for 15, 30, 60 and 90 minutes. 180 μL of supernatant was loaded into a microplate and A_340_ was determined. Points are averages of 2 samples

Sulfo-NHS-LC biotin covalently modifies primary amines, and consequently may have an effect on all surface proteins and causing secondary effects. Therefore, we measured invertase activity in yeast that were either biotinylated or incubated in PBS to see if all surface proteins become dysfunctional upon biotinylation. Invertase activity was unaffected by Sulfo-NHS-LC biotin (Supplemental Fig. S1). Therefore, biotinylation inactivated GAPDH, but not invertase, consistent with specific modification of GAPDH rather than global perturbation of either wall structure or metabolic activities.

### Enzyme assay conditions do not permeabilize the plasma membrane

In assays of cell wall enzymes, it is important that the plasma membrane is not permeabilized, so that all of the assayed activity derives from extracellular enzyme. We therefore compared PI staining before and after assaying *S. cerevisiae* for GAPDH surface activity. There was no visible increase in the fraction of propidium iodide positive yeast after assaying yeast for cell wall GAPDH (Fig. 4). Therefore, the increase in GAPDH activity seen after 100 μM DTT treatments and recovered after biotinylation was likely due to enzymes externalized by controlled by biological processes and unlikely to be caused by plasma membrane leakage.

**Figure 4.**
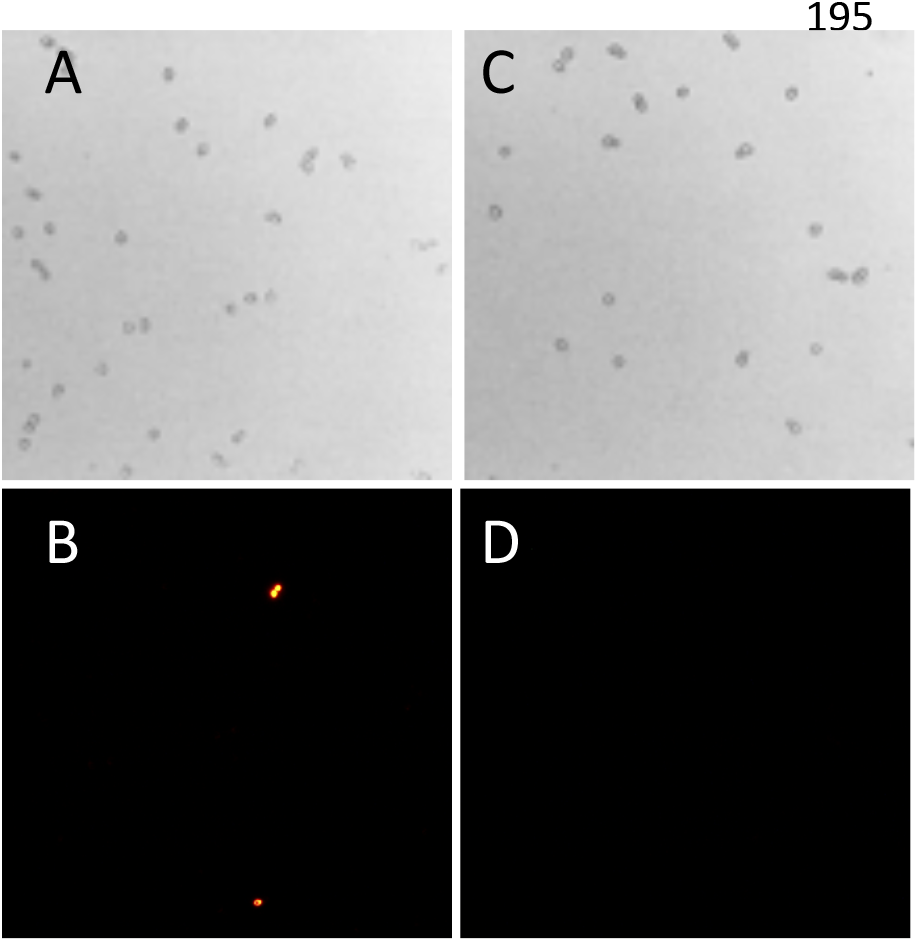
Propidium Iodide staining of *S. cerevisiae* after a GAPDH assay. *S. cerevisiae* suspended in TEA buffer pH 8.6 before (A and B) or after (C and D) assays for GAPDH surface activity. Top: brightfield image; Bottom Propidium Iodide epifluorescence in the same field.

We also found that high concentrations of reducing agents can permeabilize the plasma membrane (discussed later), so we wanted to ensure DTT concentrations used in whole cell assays for GAPDH on the surface was not permeabilizing the plasma membrane. We incubated *S. cerervisiae* in a TEA buffer at pH 8.6, at 30C in different concentrations of DTT and monitored PI fluorescence over time with flow cytometry. The results demonstrated that high concentration of DTT (1 mM or higher) permeabilized the plasma membrane, but 100 μM did not cause a significant amount of cells to become PI positive compared to a non-treated control group, and this was consistent over 90 minutes (Fig. 5). Other reducing agents including β-mercaptoethanol (5-14 mM) or TCEP (5 mM) also increased PI staining of cells, implying permeabilization of the plasma membrane (Supplemental Fig. S2). Therefore, we utilized 100 uM DTT for all subsequent assays.

**Fig. 5.**
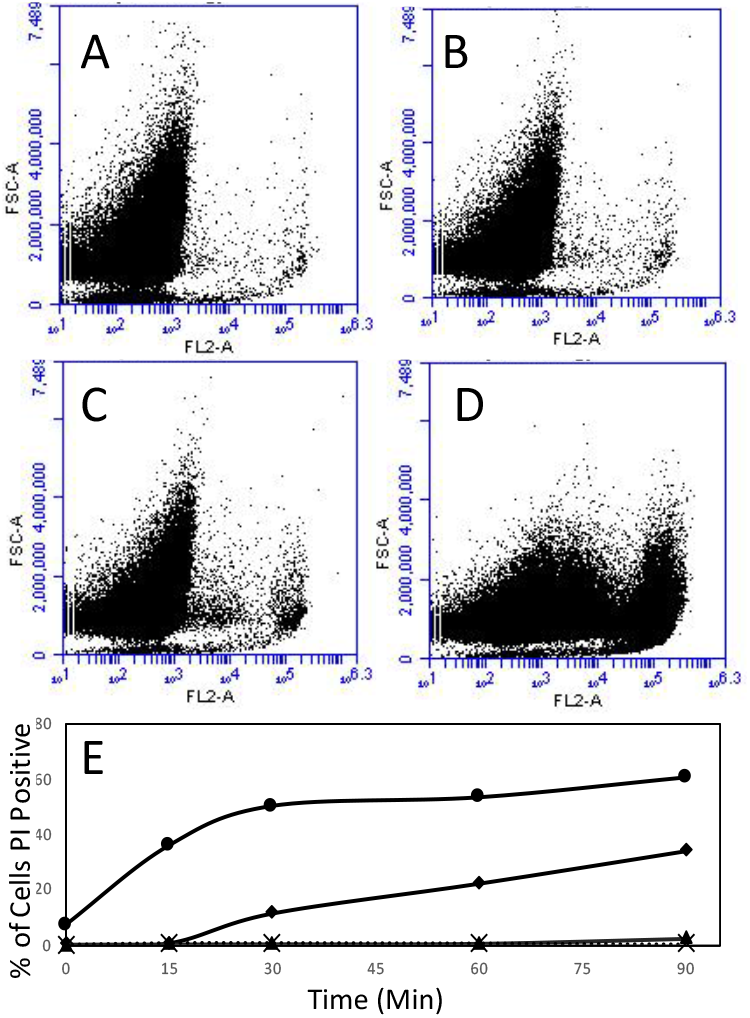
Flow cytometry of effect of DTT on Propidium Iodide staining of *S. cerevisiae* cells. **(A-D)** Propidium iodide fluorescence after 60 min incubation at 30C in TEA buffer pH 8.6. **A)** no DTT, **B)** 100 μM DTT, **C)** 1 mM DTT, **D)** 10 mM DTT. **E)** Percentage of cells which were PI positive: DTT concentrations were (X) none, (▲) 100μM, (♦) 1 mM, (●) 10 mM.

### Releasing active cell wall enzymes for *in vitro* analysis

We looked at several methods for releasing enzymatically active cell wall proteins from S. cerevisiae. It was important that the plasma membrane was not permeabilized, so that none of the activity derived from cytosolic enzyme. Since 100 μM DTT treatment did not compromise the plasma membrane, we wanted to know if we could use that concentration to extract cell wall proteins. Cells were incubated in 100 μM DTT for 60 minutes at 30C. When we used a concentration of 2.5 x 10^6^ cells per mL (the concentration used during *in situ* cell surface GAPDH assays), there was negligible GAPDH activity released into in the supernatant (not shown). However, at concentrations of 2 × 10^8^ cells per ml and above, we could monitor supernatant for NADH production. We estimate the GAPDH released by this method is less than 1% of the total GAPDH present in the wall, based on the level of activity associated with whole cells. This procedure released GAPDH when the cells were incubated at 30C, but not when they were incubated on ice (Fig. 6A). This method also released extracellular invertase from cells grown in galactose, and the yield was similarly about 1% of the total assayable invertase (Fig. 6B).

**Fig. 6.**
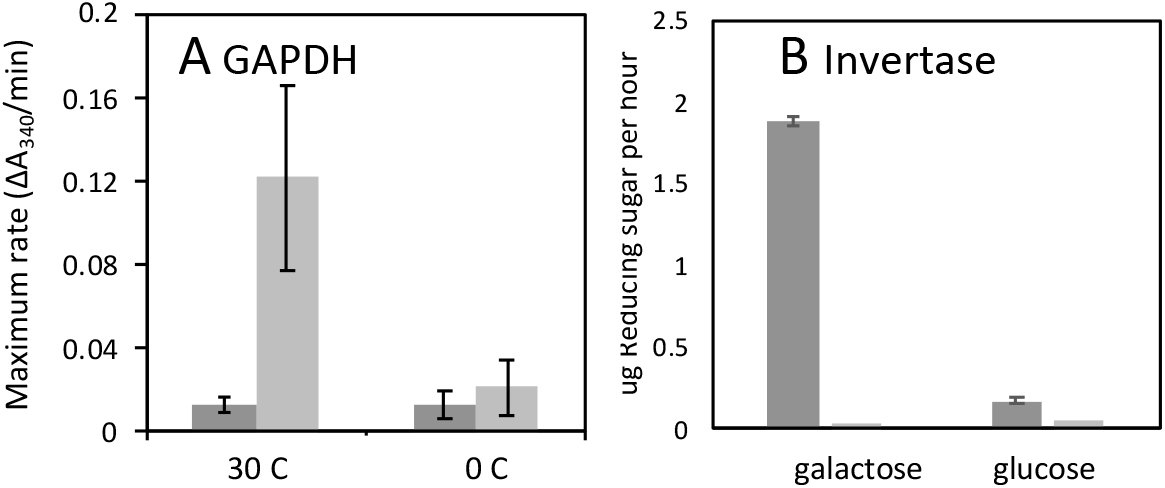
Release of cell wall enzymes from intact cells. Cells were incubated for 1 h in TEA buffer, then centrifuged and enzymes activities determined. **A)** GAPDH release after incubation at 30C or 0C in the absence of DTT (dark grey) or with 100 μM DTT present (light grey). **B)** Invertase activity after growth in galactose or glucose with added sucrose (left, dark grey), or without exogenous substrate (right, light grey).

To test other published cell wall extraction procedures, we treated cells with β1,3 glucanase, mild alkaline treatment, or reducing (5, 6, 42, 45). These methods failed to extract GAPDH without compromising the plasma membrane. Cell wall proteins extracted on ice in 100 mM Tris pH 9.4 supplemented with 2% sorbitol excludes cytosolic protein such as Cof1 (46).

However, we were unable to detect GAPDH activity in these extracts (data not shown). Treating cells with Zymolyase, a lytic β1,3 glucanase, in the presence of 1 M Sorbitol released GAPDH into the medium. However, some cells lysed rapidly, and other cells appreared to be intact but stained with PI. Therefore, neither mild base extraction nor a spheroplast procedure specifically solubilized cell wall GAPDH.

To assess effects of mild base alone, or with increased amounts of reductant, we suspended *S. cerevisiae* in 100mM carbonate buffer containing several different concentrations of β-mercaptoethanol (βME), DTT, tris(2-carboxyethyl)phosphine (TCEP), for 2 hours. Cells were pelleted by centrifugation and supernatants were assayed for GAPDH activity *in vitro* while cells were stained with propidium iodide and visualized to monitor plasma membrane leakage. At high concentrations of βME or DTT, GAPDH was released, but a large proportion of yeast treated with these concentrations readily took up propidium iodide (47) (Supplemental Fig. S2). Therefore, incubations in high concentrations of reducing agents probably released cytosolic proteins in addition to cell wall material.

## Discussion

Our results point to several practical approaches to assay of cell wall enzymes in yeast. We have screened assay procedures, and found conditions that facilitate quantitative enzyme assays without compromising the integrity of the plasma membrane. Consequently, we can estimate minimum cell surface concentrations of GAPDH as the amount of active enzyme. Additionally, selective inactivation of GAPDH, coupled with kinetics of recovery of the activity yielded a minimum estimate of the secretion rate. Thus, the results establish criteria for determination of concentrations and secretion rates for fungal cell wall enzymes.

### Cell wall GAPDH

Enzymological data leads to estimates for the amount of active enzyme on each cell surface. GAPDH enzyme assays showed NADH reduction of about 0.25 A_340_ units per hour for 5 x 10^5^ cells. Because the molar extinction coefficient of NADH is 6.2 × 10^3^ M^-1^ cm^-1^, this corresponds to production of about 3 × 10^-10^ μmol of NADH per cell per minute. Given the specific activity of yeast GAPDH, this amount of activity would result from about 4 × 10^4^ molecules of GAPDH per cell (38). This number is similar to that of other cell surface molecules such as the *S. cerevisiae* sexual agglutinins (48). Note however, that this concentration does not account for any surface GAPDH that is enzymatically inactive. For comparison, invertase, a conventionally secreted highly-expressed surface enzyme is about 100-fold higher in fully de-repressed cells (49). Therefore, cell surface GAPDH concentrations are commensurate with its frequent detection in wall proteomics studies, but are significantly lower than maximal levels of a highly expressed surface enzyme.

### Biotinylation as a tool for selectively deactivating GAPDH

Biotinylation is frequently used as a mechanism of tagging cell wall proteins for western blot analysis or proteomics (50), including unconventionally secreted proteins such as enolase (51) and the Hsp70 members Ssa1 and Ssa2 (52, 53). To our knowledge, it has not been used to deactivate enzymes *in situ*. Biotinylation ablated GAPDH activity, but did not alter external invertase activity, so not all external enzymes can be deactivated in this manner. Therefore, labeling intact cells with sulfo-NHS-LC biotin did not globally alter classical secretion and s minimally invasive. Sulfo-NHS-LC-biotin contains a charged sulfonate group, making it membrane impermeant (44). Therefore, the reagent specifically deactivated GAPDH that was external to the membrane. Propidium iodide staining and flow cytometry experiments demonstrated that the plasma membrane remained intact as GAPDH activity returned to the surface within 30-60 minutes (Figs. 3–5). Therefore, we conclude that cell surface GAPDH is specifically inactivated by sulfo-NHS-LC-Biotin, and that plasma membrane remains intact both after inactivation, and during extended incubations in assay buffer.

Sulfo-NHS-LC biotin reacts with primary amines. Based on the structure of yeast Tdh3, which is reported to be the major form of GAPDH in the cell wall with Tdh2 during exponential growth phase (1), we identified lysine residues near the catalytic cysteine, the glyceraldehyde-3-phosphate binding domain, and the NAD binding domain (54). Lysines located near the active site, include LYS160, LYS217 at 5 and 8 angstroms, as well as LYS 213, 225, 307 within 10 angstroms (Fig. 7). Therefore, it is likely that Sulfo-NHS-LC is directly inactivating GAPDH by covalently modifying one or more lysines near its active site.

**Fig. 7.**
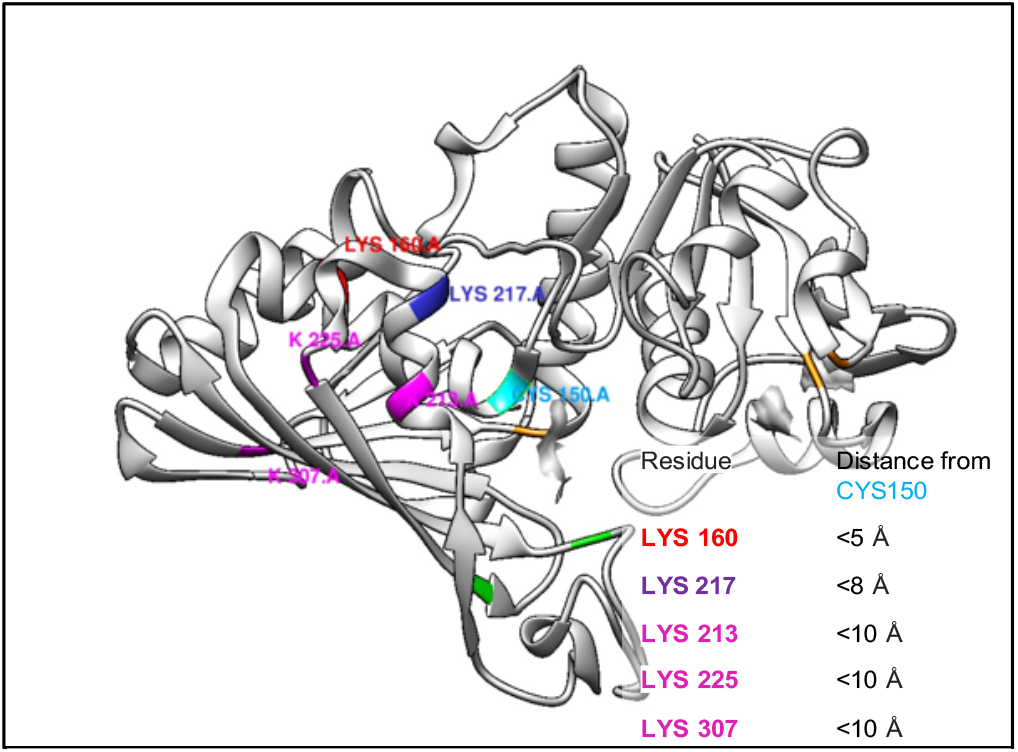
Yeast GAPDH monomer (RCSB file 4IQ8) with I active site Cys150 colored cyan, residues that bind G-3-P are colored green. Lysines are colored by distance from Cys150 (cyan), as shown in the inset table.

### Secretion of GAPDH

After inactivation of surface GAPDH, steady-state levels of activity were re-established in the wall within an hour of incubation at 30°C (Fig. 3). Thus, the rate of GAPDH secretion was about 4 × 10^4^ molecules of active enzyme per hour under these conditions. The recovery was dependent on incubation temperature, implying that the recovery was due to a secretory event rather than passive leakage from the cytosol though damaged membranes.

### Reducing agents can compromise the plasma membrane

One striking observation we made is that mM concentrations of reducing agents used to extract cell wall proteins (5, 30, 55, 56) can compromise the plasma membrane, leading to propidium iodide uptake. This is consistent with observations of Curvin et al, which used cofilin as a marker for cytosolic leakage (46).

To extract cell wall proteins for enzymology while avoiding cytosolic contamination, we recommend incubating S. cerevisiae in 100 μM DTT as described above and monitoring yeast for cytosolic permeability with propidium iodide. GAPDH and invertase are considered periplasmic (held in place between the wall and the plasma membrane) (1, 57, 58), so this technique can extract proteins associated with the innermost layer of the cell wall. We also recommend passing supernatant through a .22 μm filter to avoid contaminating extracts with unpelleted cells. Unfortunately, extraction with 100 uM DTT is inefficient, based on the observation that about 1% of the activity of GAPDH or invertase is released into the medium.

Thus, there are techniques for assay of cell wall enzymes *in situ*. A minimum estimate is 4 × 10^4^ molecules of GAPDH in the wall of each cell in exponentially growing cultures. This same amount can be secreted in an hour. We conclude that enzymatic assays are suitable for studying secretion, including unconventional secretion. GAPDH is a suitable enzyme for studying unconventional secretion. Additionally, external GAPDH can be deactivated to monitor secretion rates, and it can be readily extracted from the wall without cytosolic contamination.

## Methods

### Determining GAPDH activity at the cell surface

*S. cerevisiae* strain BY4743 was grown in YPD to log phase, pelleted, and resuspended in either 20mM sodium citrate buffer pH 5 or TBS pH 7 to an OD_600nm_ of 1.25. At this concentration 20 μL contained 5 x 10^5^ cells. 20 μL of cells from each concentration was loaded into a microfuge tube and placed on ice. To initiate the reaction, 180 μL of TEA buffer (40mM triethanolamine, Sigma, 50 mM Na2HPO4, 7.5 mM EDTA, pH 8.6) also containing 100 μM DTT, 1 mM NAD+ (Alfa Aesar) and 7 μL of 100 mg/mL glyceraldehyde-3-phosphate (Cayman Scientific or Sigma Aldrich) from frozen stocks. The cell suspension was incubated at 30C for 30 min, placed on ice for 5 minutes to retard the reaction, and then *S. cerevisiae* was pelleted by centrifuging at full speed (13,000 X G) for 1 minute. 180 μL of the supernatant was collected and A_340_ was measured on a Biotek Synergy 2 plate reader. 180 μL of supernatant from a negative control reaction of 5 x 10^5 cells without glyceraldehyde-3-phosphate or without cells were used as a blank. To determine kinetics of NADH production, cells were incubated for 0-120 minutes before analysis of supernatants.

To determine how incubation in DTT alters GAPDH activity on the surface over time, 500K yeast cells in 20 μL of TBS were mixed with in 160 μL of TEA buffer with or without DTT (100 uM) and incubated for the times stated. After incubation, NAD and glyceraldehyde-3-phosphate were added, and 2 μL of 10 mM of DTT was added to reactions lacking DTT. The tubes were incubated at 30C for 30 minutes, the supernatant was collected and analyzed for NADH production by reading an A340.

### Extraction of cytoplasmic GAPDH

*S. cerevisiae* were lysed with glass beads in PBS, with a 1:1000 dilution of yeast protease inhibitor cocktail set IV (Calbiochem), the lysate was cleared by centrifugation at 4C at full speed on a microcentrifuge, and supernatant was analyzed for GAPDH activity.

### In vitro GAPDH kinetics

10 μL of either a cell wall extract, whole cell lysate, or 10-fold dilutions were loaded into a microplate. A Biotek synergy 2 plate reader was prewarmed to 30C, 90 μL of TEA buffer containing 1 mM NAD+, glyceraldehyde 3 phosphate, and 100 μM DTT was added and an OD340 was monitored over 60 minutes. Negative control wells contained 10 μL of the buffer used to extract protein mixed with the other reagents, or extract was mixed with TEA buffer containing all of the reagents except for glyceraldehyde-3-phosphate. To calculate GAPDH activity, we determined the slope of the steepest linear part of the OD340 curve during the first 5-60 minutes.

### Biotinylation of GAPDH

Cytosolic lysate was covalently modified with or without 1mg/mL sulfo-NHS-LC biotin (ApexBio) for 1 hour. The biotinylated and non-biotinylated lysates were then washed in a 10kDa membrane cutoff filter (Sigma) with PBS, 10 μL was loaded into a microplate with 90 μL substrates and analyzed for GAPDH activity.

To biotinylate intact yeast, *S. cerevisiae* were washed and resuspended at an OD_600nm_ = 2.5 to 5 in PBS, with or without 1mg/mL Sulfo-NHS-LC biotin for 1 hour at 4C or on ice. The treated cells were washed twice and resuspended in TBS to measure GAPDH activity as above, or in citrate buffer to measure invertase.

### Whole cell Invertase assays

*S. cerevisiae* BY4741 and BY4743 were grown to an OD_600_ of 0.45-0.55 in Yeast Extract-Peptone medium with 2% galactose as carbon sourse (YPGal), concentrated to an OD_600nm_ = 1 in 20mM sodium citrate buffer (pH 5). 150 μL of this cell suspension was mixed with 50 μL of 0.4M sucrose to a final concentration of OD_600nm_ = 0.75 and 0.1M sucrose, and incubated at 30C. After ½ hour suspensions were pelleted and reducing sugar released was quantified by boiling 1:1000 dilution in tetrazolium blue (Sigma) and boiled for 3 minutes, and an OD_670_ measured in either a Spectronic 600 or Biotek Synergy plate reader. The OD_670_ was used to quantify reducing sugar against a set of glucose standards (59). All assays presented were carried out in duplicate, and are representative of 3 or more independent experiments.

To measure invertase extracted from cell walls, *S. cerevisiae* was grown to an OD_600nm_ of 0.5 in YPD (to suppress invertase) or YPGal (to derepress invertase expression), and resuspended to an of OD_600nm_ = 20, and 23, respectively, in TEA buffer (40mM triethanolamine, Sigma, 50 mM Na2HPO4, 7.5 mM EDTA, pH 8.6) containing 100 μM DTT for 60 minutes at 30C. 150 μL of 1000 x g supernatant was collected and mixed with 50 μL of sucrose in citrate buffer as stated above, except reactions were run for 60 minutes and the reaction was terminated by immediately diluting in tetrazolium blue, which is prepared in NaOH and will denature enzyme. Micrograms of reducing sugar released by invertase was measured as an A_670nm_ and compared to a glucose curve (59).

### Propidium iodide staining

*S. cerevisiae* were treated as stated, stained with either 2 to 20 ug/mL of propidium iodide (PI) (Sigma), concentrations within ranges reported for live/dead staining (4, 60, 61) and visualized under fluorescence microscopy using a TRITC filter.

### Flow cytometry

We incubated BY4743 at at a concentration of 2.5 x 10^5 per mL in TEA buffer (pH 8.6) with 0-10 mM DTT at 30C for 0-90 minutes, and at each timepoint, removed 100 uL, added PI to a final concentration of 2 ug/mL, incubated for an additional 5 minutes to ensure all dead cells take up the dye (61), and measured PI fluorescence on a BD Accuri flow cytometer.

### Cell wall extraction procedures

To generate spheroplasts, *S. cerevisiae* strain BY4743 was resuspended in PBS with or without 1M sorbitol. 1 unit of Zymolyase (Zymogen) was added to the mixture and lysis was monitored visually in the tube lacking sorbitol. Spheroplasted yeasts were identified using phase contrast microscopy at 400X magnification. The spheroplasts stabilized in sorbitol were pelleted at 2000 RPM, and supernatant was collected and assayed for enzyme activity *in vitro*. The spheroplasts were washed in PBS + 1M sorbitol, stained with PI as above (the volume of PI added did not exceed 1% of the total volume). Reducing agents for GAPDH release and cell viability were added to 2×10^6^ cells/mL in 100mM carbonate buffer containing the indicated concentrations of reducing agents at 30C for 2 hours. An aliquot of cells was stained with PI as above, remaining cells were pelleted and 10 μL of serial dilutions were used to measure GAPDH activity in the supernatant.

To extract cell wall proteins using 100 μM DTT, *S. cerevisiae* were washed 2X in TEA buffer and concentrated to an OD_600_ = 10-30. DTT was added to a final concentration of 100 μM from a 100mM frozen stock solution, the cells were incubated on either ice or at 30C for 60 minutes and then pelleted. 90% of the Supernatant was collected to avoid disturbing the pellet. In later experiments the supernatant was passed through a .22um Durapore filter (Sigma) to remove any remaining cells.

## Supplemental Data

**Supplemental Fig. S1.**
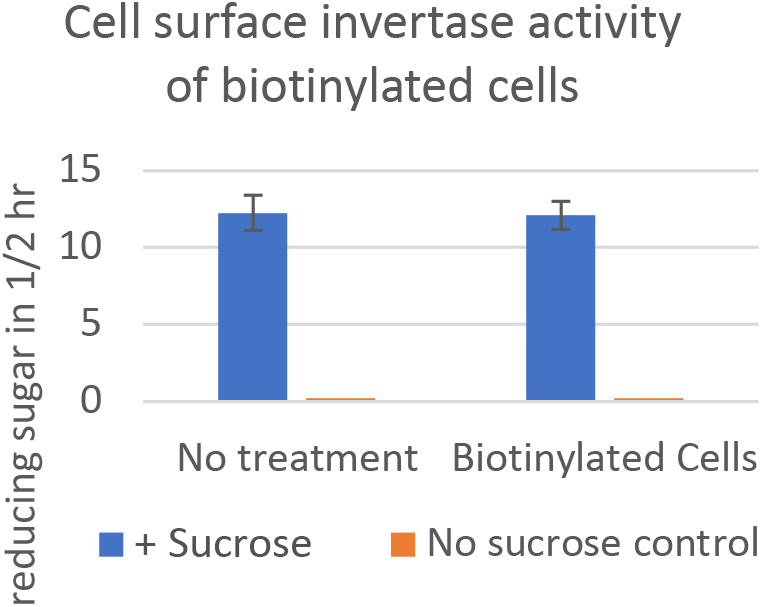
Invertase activity of intact cells with and without treatment with Sulfo-NHS-LC biotin. Yeast were grown in YPGal medium, washed, treated or not with sulfo-NHS-LC biotin, then incubated with or without sucrose, and reducing sugar determined.

**Supplemental Fig. S2.**
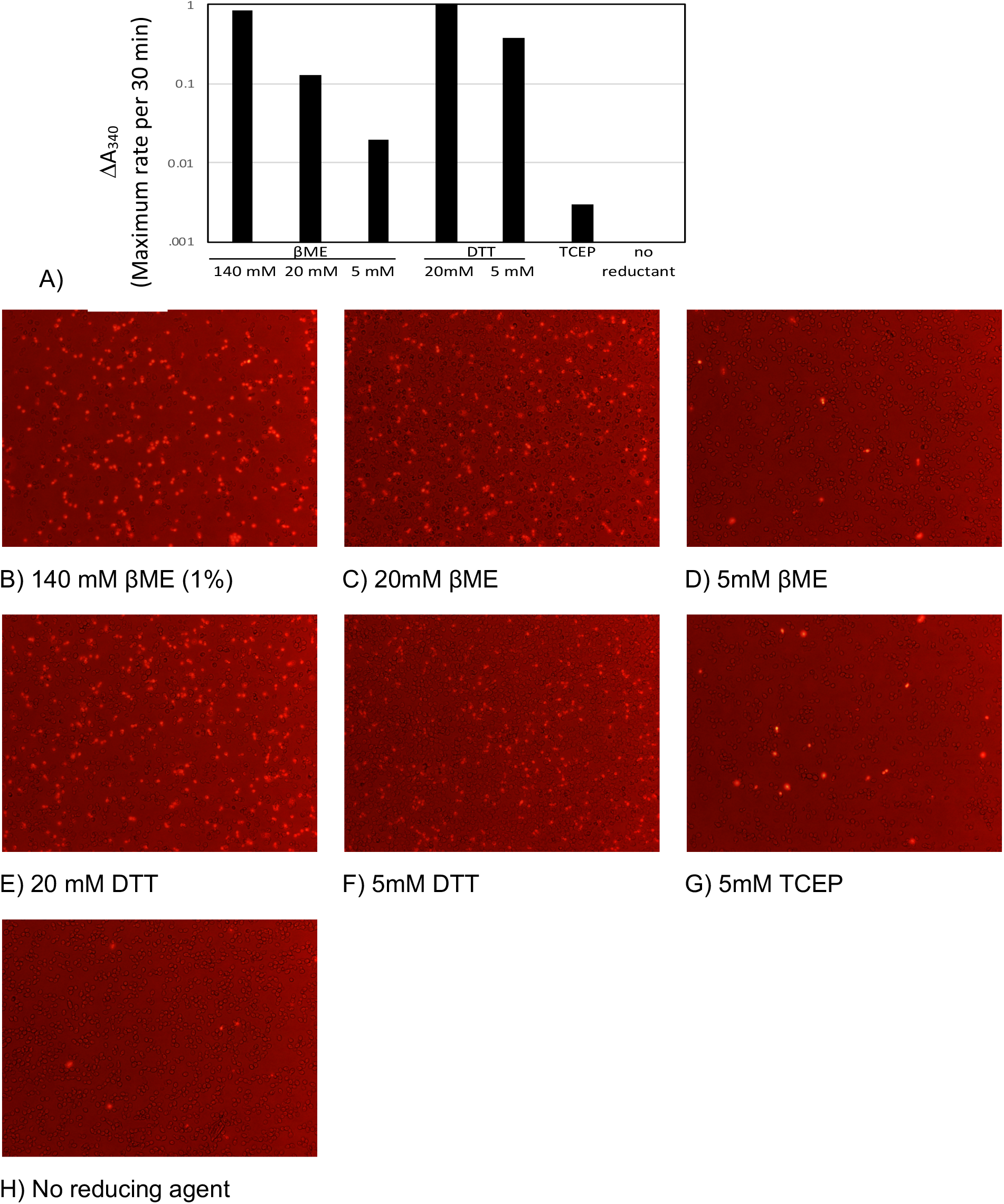
Exposing yeast to reducing agents can release cell wall enzymes, but also permeabilizes cells. **A)** GAPDH activity in supernatant from BY4743 *S. cerevisiae* after incubating in various reducing agents for 120 minutes. **B-G)** Yeast incubated in reducing agents after 2 hrs have compromised plasma membranes PI fluorescent (bright red) yeast combined with a brightfield microscopy to visualize total amount of yeast (dark spots) after exposing yeast to reducing agents compared to yeast in carbonate buffer only **(H)**. Values in A are single technical replicates and representative of 2 experiments.

